# The SV40 virus enhancer functions as a somatic hypermutation-targeting element with potential oncogenic activity

**DOI:** 10.1101/2024.01.09.574829

**Authors:** Filip Šenigl, Anni Soikkeli, Salomé Prost, David G. Schatz, Martina Slavková, Jiří Hejnar, Jukka Alinikula

## Abstract

Simian virus 40 (SV40) is a monkey virus associated with several types of human cancers. SV40 is most frequently detected in mesotheliomas, brain and bone tumors and lymphomas, but the mechanism for SV40 tumorigenesis in humans is not clear. SV40 relative Merkel cell polyomavirus (MCPyV) causes Merkel cell carcinoma (MCC) in humans by expressing truncated large tumor antigen (LT) caused by APOBEC cytidine deaminase family enzymes induced mutations. AID (activation-induced cytidine deaminase), a member of the APOBEC family, is the initiator of the antibody diversification process known as somatic hypermutation (SHM) and its aberrant expression and targeting is a frequent source of lymphomagenesis. In this study, we investigated whether AID-induced mutations could cause truncation of SV40 LT. We demonstrate that the SV40 enhancer has strong SHM targeting activity in several cell types and that AID-induced mutations accumulate to SV40 LT in B cells and kidney cells and cause truncated LT expression in B cells. Our results argue that the ability of the SV40 enhancer to target SHM to LT is a potential source of LT truncation events in various cell types that could contribute to carcinogenesis.

## Introduction

Simian Virus 40 (SV40) is a member of polyomavirus family out of which Merkel cell polyomavirus (MCPyV), BK polyomavirus (BKPyV), JC polyomavirus (JCPyV) and trichodysplasia spinulosa associated polyomavirus (TSPyV) are known to cause disease in humans (1). SV40 naturally infects monkeys and became widely known as a contaminant of Polio vaccine in 1950s-60s (reviewed in (2)). It was then discovered that SV40 transforms rodent cells efficiently and can transform cultured human cells, which led to a fear of a cancer outburst due to contaminated vaccine. This outburst never happened, but SV40 is strongly suspected of contributing to cancer formation in humans (3) although the mechanism remains unknown. SV40 is found in numerous human malignancies including mesotheliomas, brain cancers, bone tumors, and in non-Hodgkin lymphomas (4–8).

SV40 Large Tumor Antigen (LT) can transform cells by binding to retinoblastoma (RB) and p53 proteins, which interferes with cell cycle regulation and apoptosis (9–11). In MCPyV induced Merkel cell carcinoma (MCC) mutations of large T antigen (LT) causing LT truncation have a major role in carcinogenesis (12–14). Truncated forms of large T antigen with a disrupted C-terminal helicase domain are unable to bind to the viral origin of replication and initiate viral replication. Efficient replication of both MCPyV and SV40 viruses have cytotoxic effects in human cells and therefore disruption of viral replication is crucial for malignant transformation (12,15–17).

Mutations also accumulate in the SV40 *LT* C-terminus (18–20) and truncated SV40 LT retains its capability to transform cells, even when its p53 interaction domain is lost (21). While the significance of LT truncation events has not been demonstrated for SV40, it is probable that also truncated SV40 LT triggers less DNA damage response and is less immunogenic than full-length LT which is beneficial for virus persistence in host cells and might give the cells a transformative advantage similarly to truncated MCPyV LT (12,13).

Somatic hypermutation (SHM) is a process that introduces point mutations into the variable region of Ig loci and is necessary for fine modification of antibody specificity and production of high-affinity antibodies (22–24). DNA subjected to SHM is deaminated at cytidines by activation-induced cytidine deaminase (AID). The resulting deoxyuridine is recognized and processed by error-prone base excision repair and mismatch repair resulting in mutation not only at the original site of deamination but also at flanking residues (23). Little is known about the mechanism of SHM targeting. Ig loci were found to contain “mutation enhancer elements” having the ability to increase SHM of a nearby gene by two orders of magnitude or more (25,26). The SHM-targeting activity of these elements is compromised by disruption of a number of transcription factor binding sites (TFBSs), although in most cases no single binding site was critical for activity indicating their both cooperative and redundant roles (26). All known Ig mutation enhancer elements contain E-boxes (binding sites for E proteins that have important roles in B and T cell development) that were found to be important but not sufficient elements of mutation enhancers (27). Furthermore, the mutation enhancer activity of Ig enhancers is conserved through avian and mammalian species (26). SHM is also detected at a subset of non-Ig genes, both in human B cells tumors and normal germinal center B cells (28–31). Recently, we identified a number of non-Ig enhancers in the human genome with SHM-targeting activity (32). In addition, we recently showed weak somatic hypermutation (SHM) targeting activity in a MCPyV non-coding regulatory region although mutations in LT were likely caused by APOBEC3 enzymes rather than AID (33). These findings indicate that SHM-targeting activity is not an exclusive property of Ig enhancers and is governed by more general features. This finding is of particular relevance given that SHM targeting to non-Ig genes, especially when targeted to proto-oncogenes or tumor suppressors, is a known source of lymphomagenesis (29,34).

SV40 can infect several human cell types, including B cells (4,35,36) and SV40 is frequently associated with B cell derived non-Hodgkin’s lymphoma (6,8,37–40), with one study finding 62 out of 91 lymphomas to be SV40-positive (7). Strong association of SV40 with lymphomas together with frequent mutation and truncation of SV40 LT indicates that SHM in antigen-activated B cells may potentiate SV40 oncogenicity.

SV40 is also found frequently in non-B cell tumors and AID expression can be induced by inflammatory signals in non-B cells and has been detected in some non-B cell malignancies (41,42). In addition, infection with hepatitis C virus was shown to increase the rate of SHM in B cells and hepatocellular carcinoma cells (43). Together, these data indicate a potential role for SHM in SV40-mediated oncogenesis. Given the requirement of a targeting element for efficient mutation of a transcribed region and the association of SHM-targeting activity with enhancer elements, we focused on the SV40 enhancer element to test its capacity to target SHM to an adjacent LT region and generate the truncated form of LT found in human malignancies.

Here, we show that the SV40 enhancer has SHM targeting activity and that this activity can result in mutation of SV40 LT in B and non-B cancer cells. A subset of these mutations forms truncated LTs that could contribute to malignant transformation of human cells. We propose that this mutation targeting activity contributes to carcinogenesis of B cell-derived as well as non-B cell-derived malignances.

## Results

### SV40 enhancer has strong SHM targeting activity in DT40 and Ramos B cell lines

SHM is targeted to Ig genes by their enhancers and enhancer-like sequences during antibody affinity maturation (26). The mechanism of targeting is yet to be elucidated but is thought to involve modulation of RNA Pol II progression (44). The SV40 enhancer has a number of similarities with Ig enhancers (Figure 1A and 1B). It contains octamer and E-box sequences and IRF, Pu.1, and NF-kB bind sites, all known to be important for SHM targeting activity in B cells (26). We measured the SHM targeting activity of the SV40 enhancer using established GFP fluorescence loss assays based on reporter vectors expressing GFP (GFP2 assay) (25) or a hypermutation target sequence (HTS)-GFP fusion gene (GFP4 and GFP7 assays) (26,32). HTS contains numerous SHM hotspot motifs designed to yield stop codons upon mutation of cytidine, allowing the vector to report SHM activity with high sensitivity by virtue of the loss of GFP fluorescence (Figure S1). These assays have been used to identify and characterize both Ig and non-Ig SHM-targeting enhancers as well as for mapping SHM-susceptible regions in the human B cell genome (25–27,32). An important feature of these reporter vectors is that GFP expression is driven by a strong, constitutively active promoter and addition of a strong enhancer increases GFP transcription and fluorescence levels modestly or, in some cases, not at all —thereby allowing effects on SHM to be distinguished from those on transcription levels (26,32).

**Figure 1.**
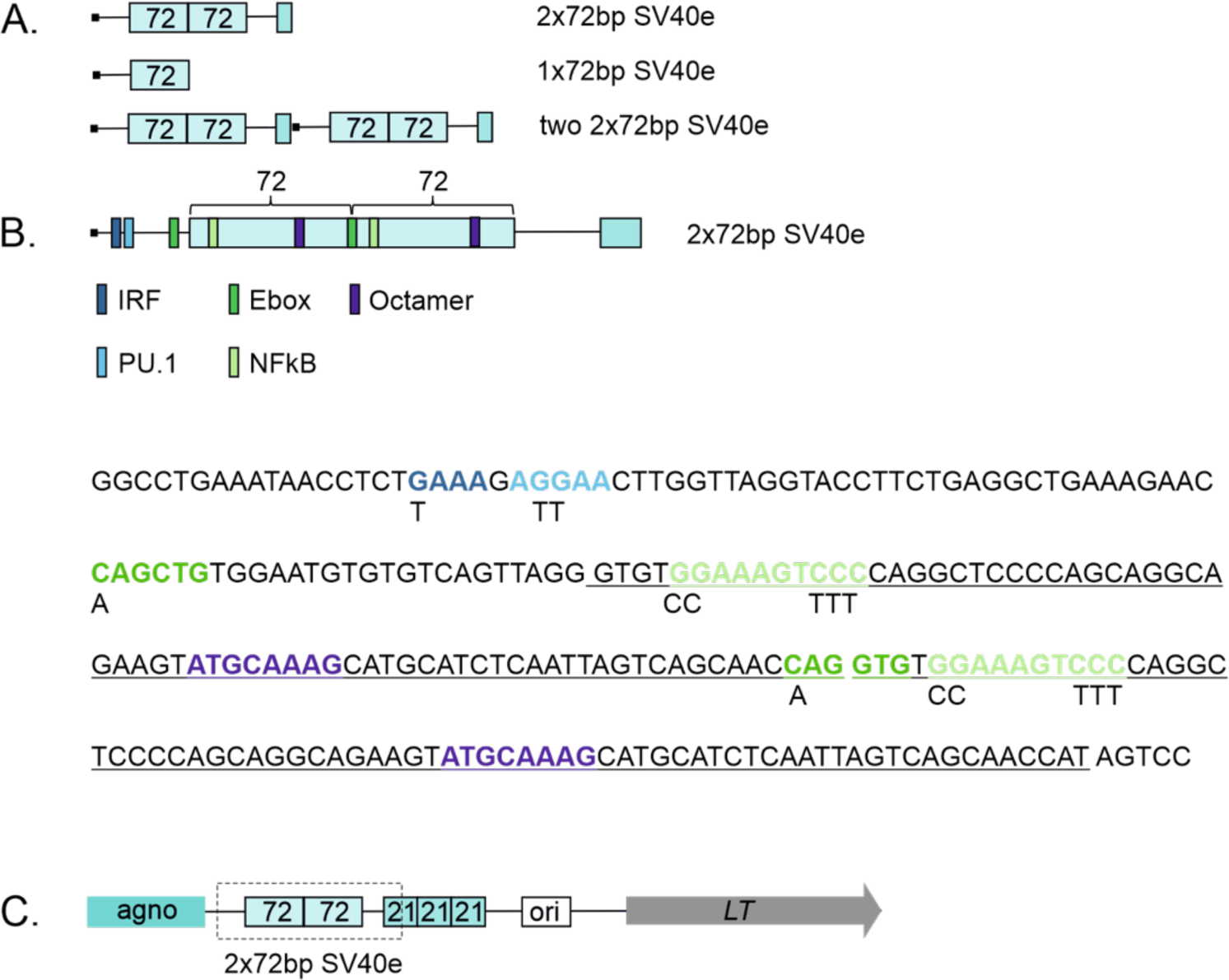
Schematic figures of SV40 enhancer and genome inserted *LT* area. A. SV40 enhancer region and its modifications measured in GFP fluorescence loss assay. B. Detailed representation and sequence of SV40 enhancer. Locations TFBSs are indicated in different colors. 72bp repeats are underlined in sequence. Changed nucleotides in mutated TFBSs are shown under the original sequence. C. SV40 *LT* expression cassette integrated to Ramos, UO-31 and DT40 genome. Representative locations of agno-protein (agno), 72bp and 21bp-repeat regions, origin of replication (ORI) and *LT* area are shown. Dashed line rectangle indicates region measured in GFP fluorescence loss assay.

We found that the SV40 enhancer is a strong mutation enhancer in the chicken bursal lymphoma cell line DT40, where it increases SHM-mediated GFP fluorescence loss 25-fold (GFP fluorescence loss 1.89% in empty reporter vs 47.5% with SV40 enhancer, p<0.0001, Figure 2A). This SHM targeting activity is comparable to that of the Ig heavy enhancer core sequences (GFP fluorescence loss 20.2%, p<0.0001 compared to empty reporter, Figure 2A). The SV40 enhancer also increases SHM in the human Burkitt’s lymphoma cell line Ramos, driving a 40-fold and 85-fold increase in GFP fluorescence loss when present in one or two copies, respectively (0.021% in empty vector vs. 0.855%, p<0.0001 and 1.755%, p<0.0001 respectively, Figure 2C), indicating that mutation enhancer activity is evolutionarily conserved in vertebrates. This is in agreement with data reporting human and mouse Ig enhancer mutation activity in chicken DT40 B cell line or chicken enhancer element mutation activity in human Ramos B cell line (26,32).

**Figure 2.**
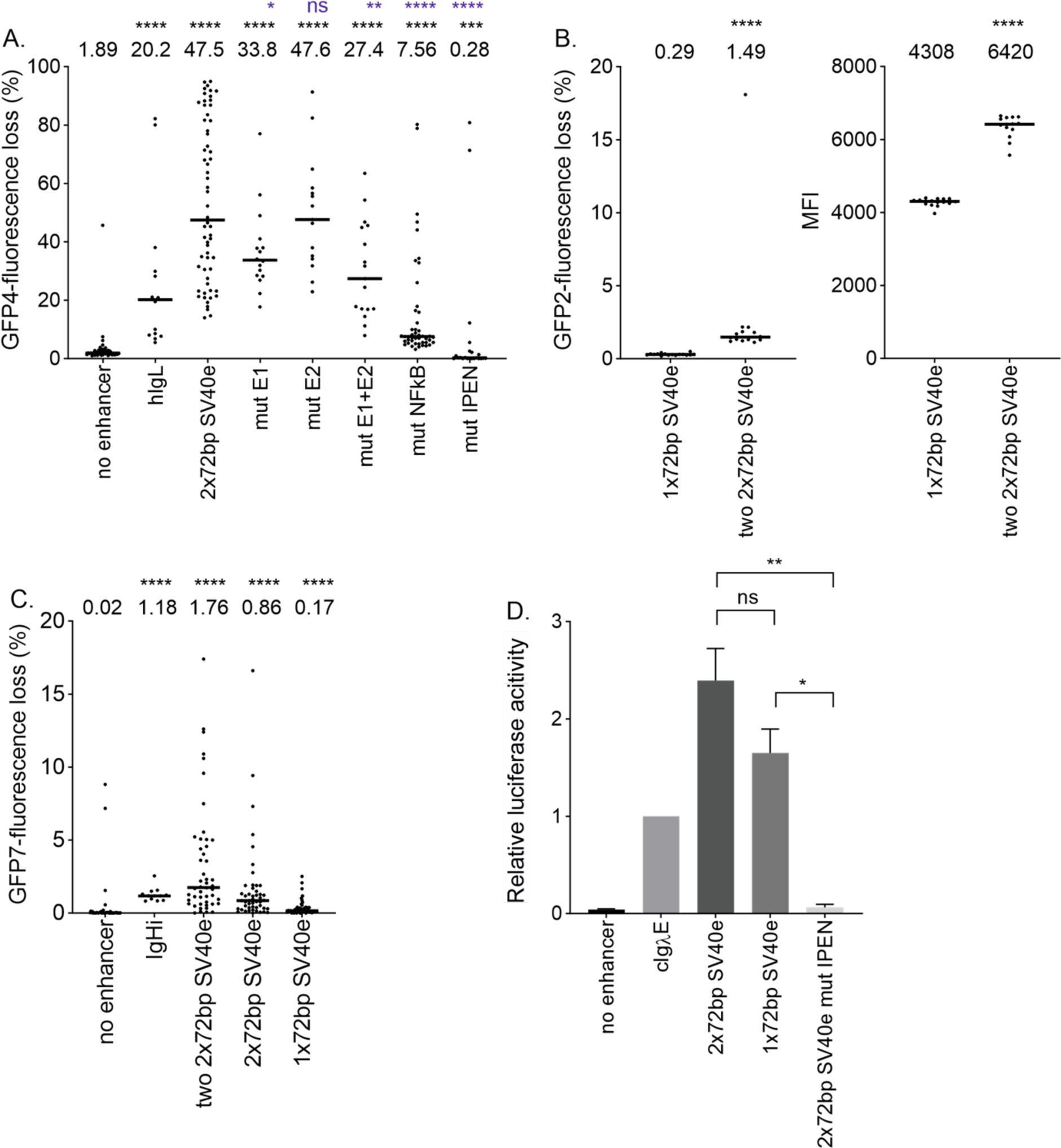
SV40 enhancers SHM targeting activity in chicken and human B cells. A. SHM targeting activity of 2×72bp SV40 enhancer and TFBS mutants measured in GFP fluorescence loss assay in GFP4 reporter in chicken DT40 cell line. Empty GFP4 reporter vector was used as negative control and human Ig lambda (hIgL) enhancer core was used as positive control. Median values are shown. Statistical significance compared to negative control is indicated in black and statistical significance compared to 2×72bp enhancer is indicated in blue. Mann-Whitney U-test was used to calculate statistical significances. Mutated TFBS binding site(s) are indicated in x-axes naming. IPEN: IRF, PU.1., E-box and NF-KB binding site mutated. B. SHM targeting activity of 1×72bp SV40 enhancer and two 2×72bp SV40 enhancer measured in GFP fluorescence loss assay in GFP2 reporter in chicken DT40 cell line and mean fluorescence intensity (MFI) of GFP resulted by 1×72bp SV40 enhancer and two 2×72bp SV40 enhancer. Median values are shown. Mann-Whitney U-test was used to calculate statistical significance. C. SHM targeting activity of two 2×72bp SV40 enhancer, 2×72bp SV40 enhancer and 1×72bp SV40 enhancer in Ramos human B cells. Empty GFP7 reporter vector was used as negative control and human Ig heavy intronic enhancer core (IgHi) was used as positive control. Median values are shown. Mann-Whitney U-test was used to calculate statistical significances. D. Enhancer activity of 2×72bp SV40 enhancer, 1×72bp SV40 enhancer and 2×72bp SV40 IPEN mutant enhancer measured in luciferase assay. Chicken Ig lambda enhancer (cIgλE) was used as positive control and empty vector as negative control. Two replicate measurements were performed. Fold changes over cIgλE are shown. Unpaired t-test was used to calculate statistical significances. **** <0.0001, *** <0.001, ** <0.01, * <0.05

To determine whether SV40 mutation enhancer activity depends on similar TFBSs compared to Ig enhancer, we created TFBS mutants where binding sites were mutated individually (E-box1, E-box2, NF-kB) or in various combinations (both E-boxes, or E-boxes together with IRF, PU.1 and NF-kB binding sites, termed IPEN) and tested them in the GFP fluorescence loss assay in DT40 cells (Figure 1B and 2A). Mutation of E-box1 decreases GFP fluorescence loss by 28.8% compared to WT enhancer (33.8% vs. 47.5%, p= 0.0423) (Figure 2A). Mutation of E-box2 has no or little effect on SV40 enhancer SHM targeting activity alone (GFP fluorescence loss 47.6% vs. 47.5%, p= 0.7605), but has a small additive effect with the mutation of the other E-box (GFP fluorescence loss 27.4% vs. 47.5%, p= 0.0012). Mutating the NF-kB site decreases SHM targeting activity by 84.1% (GFP fluorescence loss 7.56% vs. 47.5%, p <0.0001). Mutating IRF, PU.1, both E-boxes, and NF-kB binding sites abolishes SHM targeting activity completely (GFP fluorescence loss 0.28% vs. 47.5%, p <0.0001), These data indicate that the SHM-targeting activity of the SV40 enhancer is dependent on the same transcription factor binding sites as Ig enhancers (26).

The SV40 enhancer naturally contains one or two direct 72 bp sequence repeats depending on the strain of the virus (2). The number of 72 bp repeats affects virus properties such as virus growth and replication efficiency (2,3,45). To test if the number of repeats affect mutation enhancer activity, we tested the one (1×72bp) and two-repeat (2×72bp) SV40 enhancers for their SHM targeting activity using GFP fluorescence loss assay. SHM targeting activity was 5-fold decreased in 1×72bp when compared to 2×72bp enhancer in Ramos (0.17% vs. 0.86%, p<0.0001, Figure 2C). Comparison of 1×72bp enhancer with two 2×72bp SV40 enhancers (containing total of four 72bp repeats) revealed 5-fold decrease of SHM targeting activity in DT40 (0.29% vs. 1.49% p <0.0001) and 10-fold decrease in Ramos (0.17% vs. 1.76%, p <0.0001, Figure 2B). The mean GFP fluorescence intensity of cells containing a vector with two 2×72bp SV40 enhancers was 1.5-fold higher than 1×72bp SV40 enhancer in DT40 (p <0.0001) (Figure 2B). We also measured the gene expression induced by 2×72bp SV40 enhancer, 1x 72bp SV40 enhancer, and 2×72bp SV40 enhancer IPEN mutant with a luciferase assay (Figure 2D). There was no difference in fold changes between 2×72bp enhancer and 1×72bp enhancer (p=0.1256). The fold change of activity of 2×72bp and 1×72bp enhancers compared to 2×72bp enhancer IPEN mutant was significantly greater (p=0.0100 and p=0.0121, respectively).

As in previous studies (32,46), we used GFP mean fluorescence intensity to assess the effects of various enhancers on levels of transcription. In Ramos cells, the two 2×72bp SV40 enhancer increased GFP mean fluorescence intensity while the one 2×72bp SV40 enhancer and one 1×72bp SV40 enhancer did not (Figure S2). The intensity driven by a single 2×72bp SV40 enhancer was not affected by the loss of one 72bp repeat in Ramos.

AID expression in DT40 and Ramos cells was confirmed using qPCR and Western blot methods (Figure S3C, D). To confirm the dependency of the observed GFP fluorescence loss on AID activity, we performed GFP fluorescence loss experiments in AID deficient Ramos cells (Ramos AID-/-). SHM targeting activity of the SV40 enhancer was entirely dependent on AID (GFP fluorescence loss 0%; Figure S5).

Reconstitution of AID expression in the AID-deficient cells by transduction with a retroviral vector bearing an AID-mCherry expression cassette fully restored SHM and GFP fluorescence loss (Figure 3D).

**Figure 3.**
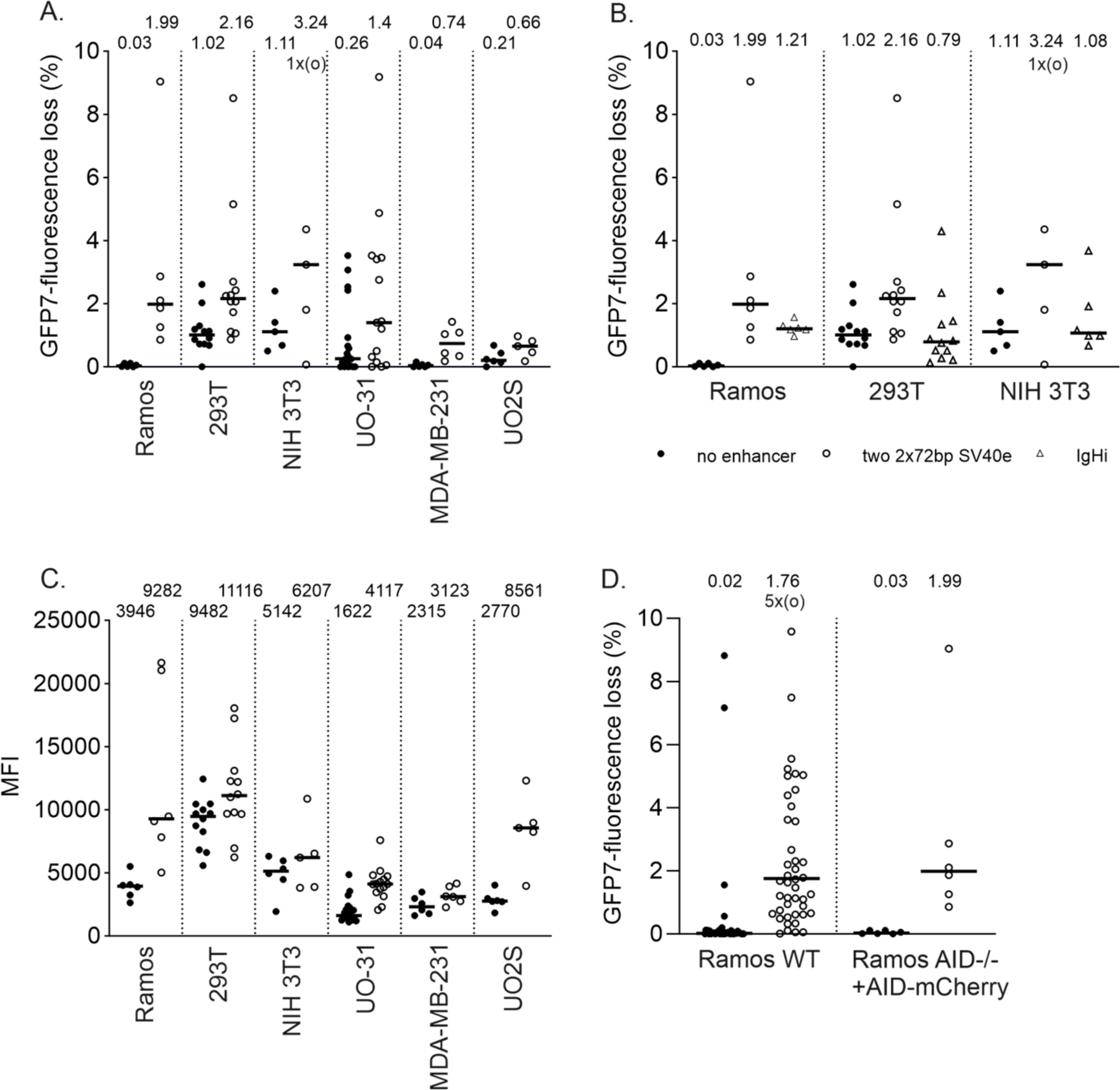
SV40 enhancers SHM targeting activity in non-B cells. A. GFP fluorescence loss of two 2×72bp SV40 enhancer measured in GFP fluorescence loss assay in GFP7 reporter in UO-31 cell line and Ramos AID-/-, 293T, NIH 3T3, MDA-MB-231 and UO2S cell lines transduced with AID-mCherry expression vector. Median values are shown. B. GFP fluorescence loss of two 2×72bp SV40 enhancer and IgHi measured in GFP fluorescence loss assay in GFP7 reporter in Ramos AID-/-, 293T, NIH 3T3 cell lines transduced with AID-mCherry expression vector. Median values are shown. C. Mean fluorescence intensities of two 2×72bp SV40 enhancer measured in GFP fluorescence loss assay in GFP7 reporter in UO-31 cell line and Ramos AID-/-, 293T, NIH 3T3, MDA-MB-231 and UO2S cell lines transduced with AID-mCherry expression vector. D. GFP fluorescence loss of two 2×72bp SV40 enhancer measured in GFP fluorescence loss assay in GFP7 reporter in Ramos WT and Ramos AID-/- cell line transduced with AID-mCherry expression vector. Median values are shown.

### SV40 enhancer has SHM targeting activity in non-B cells

SV40 can infect various cell types and is linked to non-B cell malignances such as mesotheliomas and bone and brain tumors (47). Therefore, we also investigated SV40 enhancer targeting activity in cell lines of other than B cell origin. Human renal cell carcinoma UO-31 had detectable AID mRNA expression corresponding to 13% of AID expression in Ramos (Figure S3A) measured with RT-qPCR. AID expression in UO-31 was further confirmed using immunohistochemistry and western blot (Figure S3B). In UO-31 cells, SV40 enhancer had SHM targeting activity measured as a GFP fluorescence loss in clones bearing the GFP7 SHM-reporter retroviral vector (more than 5-fold increase, Figure 3A). Sequencing of the GFP7 coding sequence revealed 16 mutations out of which 10 (62.6%) were targeted to Cs confirming the SHM targeting into the vector (supplementary Figure S4).

Due to the scarcity of non-B cell lines expressing AID, we selected four cell lines of different tissue and species origin and transduced them with the GFP7 vector: mouse embryonic fibroblast NIH3T3, human embryonic kidney epithelial cell 293T, human osteosarcoma U20S, and human breast adenocarcinoma MDA-MB-231. Analysis of clones found no significant GFP fluorescence loss, confirming that these lines do not have active SHM. Ramos AID-/- cells were used in the parallel experiment as a control cell line with inactive SHM. Only 293T cells exhibited significant percentage of GFP-negative cells which was due to low sensitivity of this cell line to blasticidin selection, which is not able to fully remove cells with transcriptionally silenced GFP7 vector (Figure S5). We then ectopically expressed AID in the five parental cell lines. 293T, NIH3T3, U20S and MDA-MB-231 were transduced with the AID-mCherry expression vector, with Ramos AID^-/-^ used as a control to verify that this vector is able to restore the pattern of GFP fluorescence loss we observed in wild-type Ramos cells. This panel of cells was then transduced with the GFP7 reporter vector lacking or bearing the SV40 enhancer. All non-B cells exhibited significant GFP fluorescence loss even in the control vector and the presence of the SV40 enhancer further increased the GFP fluorescence loss 2-20-fold (Figure 3A). GFP fluorescence loss does not correlate with increase of GFP fluorescence intensity indicating that the SHM increase is not solely caused by an increase in transcription (Figure 3C). GFP fluorescence loss in Ramos cells (Ramos AID-/- transduced with AID-mCherry expression vector) replicated the results obtained with wild-type Ramos cells with a 60-fold GFP fluorescence loss increase upon SV40 enhancer insertion. (Figure 3D). Thus, the SV40 enhancer is a versatile SHM-targeting element able to target mutations in various cell types provided they express AID. Notably, this property is distinctive to the SV40 enhancer since the well-characterized IgH intronic enhancer is a potent SHM-targeting element in B cell-derived cell lines (Ramos, DT40) but has neither SHM-targeting nor enhancer activity in non-B cells (Figure 3B, S6).

### SV40 *LT* region accumulates mutations in B cells leading to truncated LT expression

Our experiments demonstrate that the SV40 enhancer is able to target SHM into the reporter GFP7 transcription unit. To determine whether the SV40 enhancer is able to target SHM in the context of SV40 virus genome and thus modify the viral genome, we introduced the SV40 large T antigen transcription unit including the enhancer region into Ramos B cells. We also included the upstream SV40 agnoprotein coding sequence, which is transcribed in the reverse orientation from the same promoter-enhancer as *LT* (figure 1B) to mimic the natural genomic context of the SV40 enhancer. Following a 12-week culture, we sequenced the *LT* region to assay for the accumulation of mutations. We found 47 mutations out of which 26 (55.3%) were targeted at Cs (Figure 4). 19.1% of all mutations and 34.6% of C-targeting mutations were at AID hotspots (WR**<u>C</u>**, Figure 4), with C-targeting mutations being significantly enriched at hotspots (p= 0.0025). Three of the mutations introduced insertions or deletions causing frameshifts and two of the substitutions generated STOP codons (Figure 4). In parallel experiments in DT40 cells, we also detected the generation of a STOP codon in the *LT* coding sequence. Furthermore, we were able to detected an approximately 50 kDa truncated LT protein corresponding to the deletion of C at position +1727 respective to the transcription start site resulting in the generation of STOP codon at position +1728 (Figure 5). These results further confirm that SV40 enhancer-induced SHM can cause truncation events in LT in B cells.

**Figure 4.**
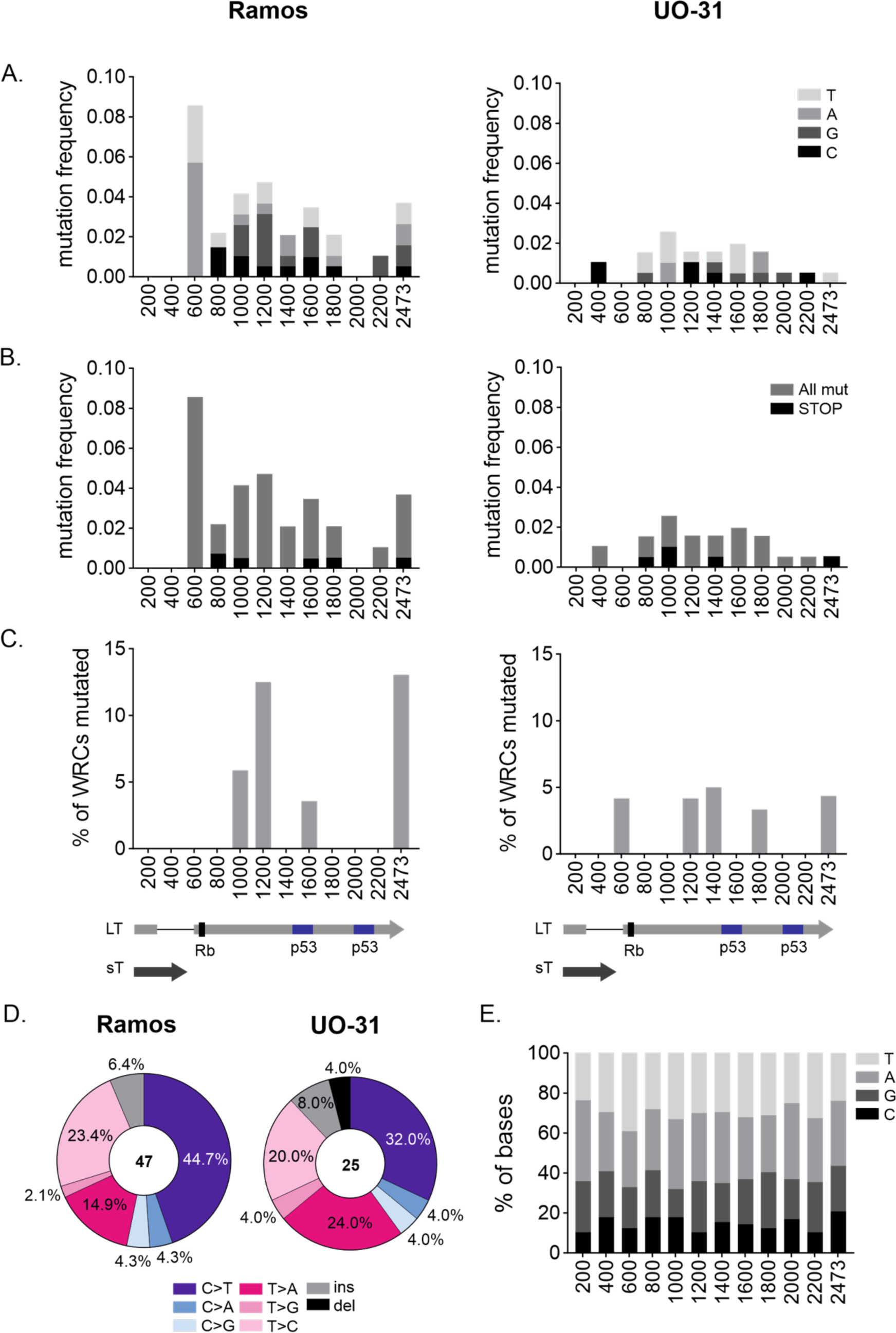
Mutation type and distribution of *LT* area in Ramos and UO-31 cells. A. Overall mutation distribution along SV40 *LT* area. Mutations targeting different bases are indicated in different colors. Mutation frequency (number of mutations/number of sequences per bin) is shown. B. Overall and STOP-codon forming mutation distribution along SV40 *LT* area. Overall and STOP-mutation are indicated in different colors. Mutation frequency (number of mutations/number of sequences per bin) is shown. C. % of mutations targeting AID hotspot WR**<u>C</u>** per BIN along SV40 *LT*. D. Mutation types in *LT* area. Substitutions are presented in six substitution classes: C>A, C>T, C>G, T>A, T>C, T>G. In addition, deletions and insertions are shown. E. Each bin of SV40 *LT* presented by its base content. For panels A-C *LT* area is divided to 200 bp bins except final bin being 273 bp. A schematic of *LT* area with locations of sT, LT, Rb1-binding site and two-part p53 binding site are depicted below the graphs.

**Figure 5.**
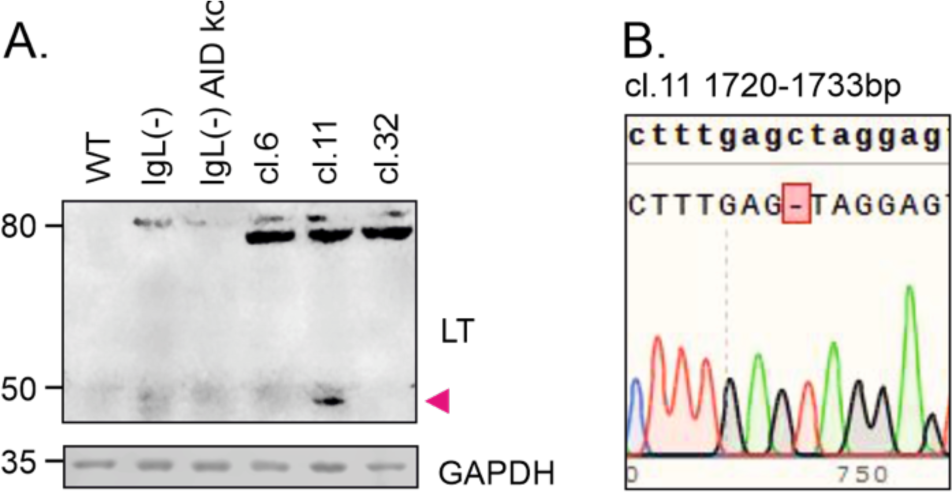
SV40 *LT* STOP codon mutation and truncated LT expression in DT40 cells. A. SV40 LT and GAPDH protein expression in DT40 cell lines measured with western blot. SV40 LT and GAPDH expression was measured from DT40 wild-type, IgL(-) and IgL(-) AID knock-out cell lines. Clones (cl.) 6, 11 and 32 are IgL(-) cells where to SV40 *LT* expression cassette is integrated to Ig locus. Protein size is indicated in left. Truncated LT is marked with arrow. B. Sequencing chromatogram from clone 11 lined with reference SV40 *LT* sequence. Deletion of C base at position 1727 results in-frame TAG STOP codon immediately after deletion.

### SHM targeting activity of SV40 enhancer induces mutations in *LT* in a non-B cell line

Since SV40 enhancer targets SHM both in B cells and in non-B cells, we introduced the SV40 transcription unit into UO-31 cells which exhibit endogenous AID expression and SHM targeting. After 12 weeks of culture, we detected 25 mutations out of which 10 (40.0 %) were targeted to Cs (Figure 4). Compared to Ramos cells the mutation load was lower, which likely reflects the lower AID expression in UO-31 cells. Six mutations were in AID hotspots (Figure 4) and 60.0% of all C targeting mutations were at WR**<u>C</u>**, indicating significant hotspot enrichment in C targeting mutations (p= 0.0012). Three of the mutations introduced insertions or deletions causing frameshifts and two of the substitutions generated STOP codons (Figure 4).

## Discussion

We characterized SV40 enhancer SHM-targeting properties in various cells types, with implications for both virus fitness and oncogenesis. We found that the SV40 enhancer has SHM targeting activity comparable to Ig enhancers in B cells and that unlike Ig enhancers, the SV40 enhancer is also active in non-B cells where it can stimulate SHM up to 20-fold. Furthermore, in cells expressing AID, the SV40 enhancer can substantially increase the probability of LT truncation mutations. LT truncation has been implicated as the event necessary for human cell malignant transformation of a related MCPyV polyomavirus (12) and has been observed with SV40 in human cell lines (48,49). Comparison of SV40 enhancer SHM-targeting activity in various cell types revealed that it is most potent in B cells, where it shows a one order of magnitude higher stimulation of SHM than in non-B cells. We found that SV40 has SHM targeting activity in human, chicken, and mouse cells, arguing for an evolutionary conserved mechanism of SV40 enhancer SHM targeting activity. However, the stronger SHM-targeting activity of the SV40 enhancer in B cells than in non-B cells might be due to the lack of B cell-specific binding factor(s) in non-B cells. This is supported by the fact that the B cell-specific IgHi enhancer exhibits no SHM targeting activity in non-B cells.

SV40 can integrate into the host genome and depending on host cell permissiveness (permissive or non-permissive), it can replicate. In semi-permissive human cells, where SV40 replicates with variable efficiencies depending on the cell type, virus replication results in host cell death. Stable infection of human cells requires restriction of virus replication efficiency. One mechanism to decrease replication efficiency is LT truncation, which is found in human SV40-transformed cell lines (48,49). This is a major difference compared to rodent cells which are non-permissive and thus do not require LT truncation events to limit virus replication and propagation. Thus it is not surprising that truncation of LT has no clear benefits but rather is a disadvantage in the transformation of rodent cells (18,50–52). In human cells, SV40 small T antigen (sT) and LT are widely used for cell transformation and in such experiments LT truncation is not required in the absence of whole SV40 genome. Thus, truncation of LT in human cells could provide a route for cellular transformation in vivo.

Our results show that the SV40 enhancer can facilitate mutation of *LT* (C-terminus/helicase domain) via SHM if cells are expressing AID. We found this to occur in B cells and at lower rate in kidney cells. Kidney cells are SV40’s natural host cells in monkeys and are considered as the SV40 reservoir in humans along with blood cells (47). Furthermore, traces of SV40 have been detected in human renal cell carcinomas (53,54). Among different cancer types, evidence for the presence of SV40 is most prevalent in B cell lymphomas (47). Moreover, among non-Hodgkin lymphomas, SV40 is most frequently found in diffuse large B cell lymphoma and follicular lymphoma, both linked to aberrant SHM (34,55).

Our results show that the same B-cell TFBSs are important for the SHM-targeting activity of the SV40 enhancer and Ig enhancers in B cells (Figure 2A) (26,27). The distinctive ability of the SV40 enhancer to target SHM in non-B cells argues that certain non-B cell factors contribute to SHM targeting in non-B cells. Such factors might include AP-2 and AP-1 transcription factors which are shown to bind SV40 enhancer (56). AP-2 and AP-1 are also expressed in kidney cells and have been implicated in early carcinogenesis of renal cell carcinomas (57–60).

AID expression is a requirement for SHM and is usually limited to B cells but can be induced in non-B cells by inflammatory signals. It is attractive to think that AID expression and SHM contribute to SV40-mediated oncogenesis given that SV40 LT is sufficient to transform cells and maintain the transformed phenotype but might not be enough to lead to full cancer progression (2), with additional factors such as inflammation being required. SV40-associated cancers are more frequent in immunocompromised individuals (56) further arguing that overcoming SV40 virus immunogenicity, either by immune system malfunction or restriction of virus replication, can contribute to the development of cancer.

Our results show that 2×72bp enhancer has stronger SHM targeting activity than 1×72bp enhancer (Figure 2). The archetype SV40 enhancer contains one 72bp repeat in its enhancer region and reference laboratory strain 776 contains two repeats (2,61). However, duplication of the 72bp repeat can occur in some cell types (62) and two 72bp repeats can significantly increase transcription of the target gene (63). In addition, 2×72bp repeat virus quickly replaces one-repeat virus in culture when co-transfected, which indicates a growth advantage of the two-repeat virus (64). Nevertheless, 1×72bp variant has been detected in tumors more often than 2×72bp (2), which may be attributable to the higher prevalence of the 1×72pb variant, unstable nature of polyomavirus regulatory region (56), or loss of one repeat by AID-mediated recombination during tumor passage (26). From the perspective of preserving the virus in the genome, 1×72bp might be favored as it limits transcription/replication and thus immunogenicity.

In rodents, the SV40 early region (enhancer and *LT*) is usually found intact and integrated into the genome without apparent integration site preference (2). Research into sites of SV40 integration in the human genome is limited, but integration has been shown at least in osteosarcomas (65) and in immortalized human fibroblasts (66). It has been proposed that polyomavirus LT expression is needed in early cell transformation but becomes dispensable later in tumor development (hit-and-run mechanism) (67,68) in which case integration is not necessary. Our study reveals the first viral enhancer exhibiting strong SHM-targeting properties and demonstrates that this activity could play a substantial role in SV40 virus oncogenic potential.

**Figure S1.**
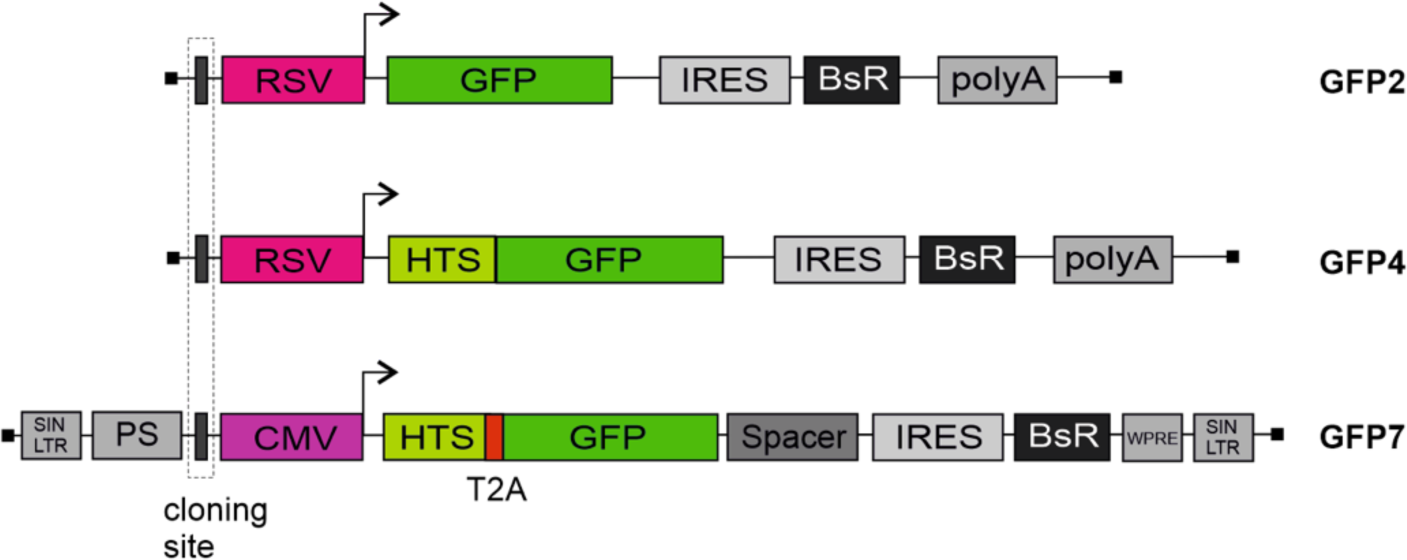
GFP reporter assay vectors. Map of GFP2, GFP4 and retroviral GFP7 SHM reporter vector. SV40 enhancer and enhancer modification cloning sites are indicated with dashed line rectangle. SpeI/NheI cloning site was used in GFP2 and GFP4 vectors. HpaI cloning site was used in GFP7 vector. Transcription direction is indicated with arrow. RSV, Rous sarcoma virus promoter; CMV, Cytomegalovirus promoter; HTS, hypermutation targeting sequence; GFP, green fluorescent protein; T2A, self-cleaving T2A peptide; IRES, internal ribosome entry site; BsR, blasticidin resistance; polyA, SV40 polyA signal; PS, packaging signal; SIN LTR, self-in-activating long terminal repeat; Spacer, sequence that place Bsr outside of the SHM target window; WPRE, woodchuck hepatitis virus posttranscriptional regulatory element.

**Figure S2.**
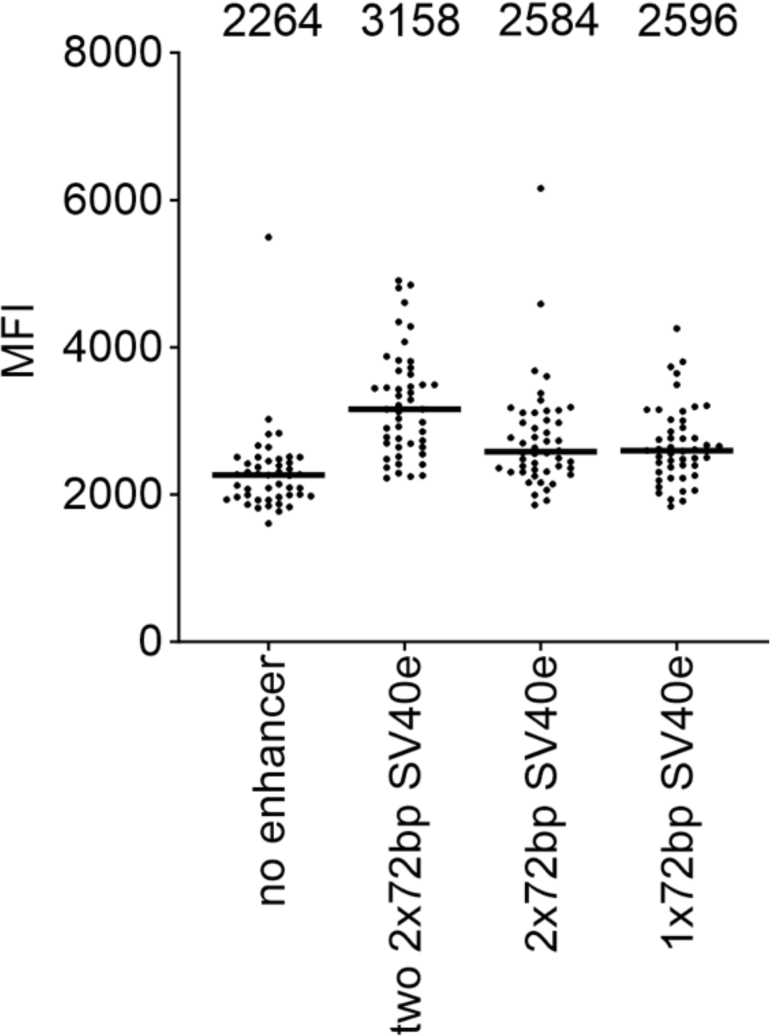
Mean fluorescence intensity of SV40 enhancers. Mean fluorescence intensity of two 2×72bp SV40 enhancer, 2×72bp SV40 enhancer and 1×72bp SV40 enhancer measured in GFP fluorescence loss assay in GFP7 reporter in Ramos human B cells.

**Figure S3.**
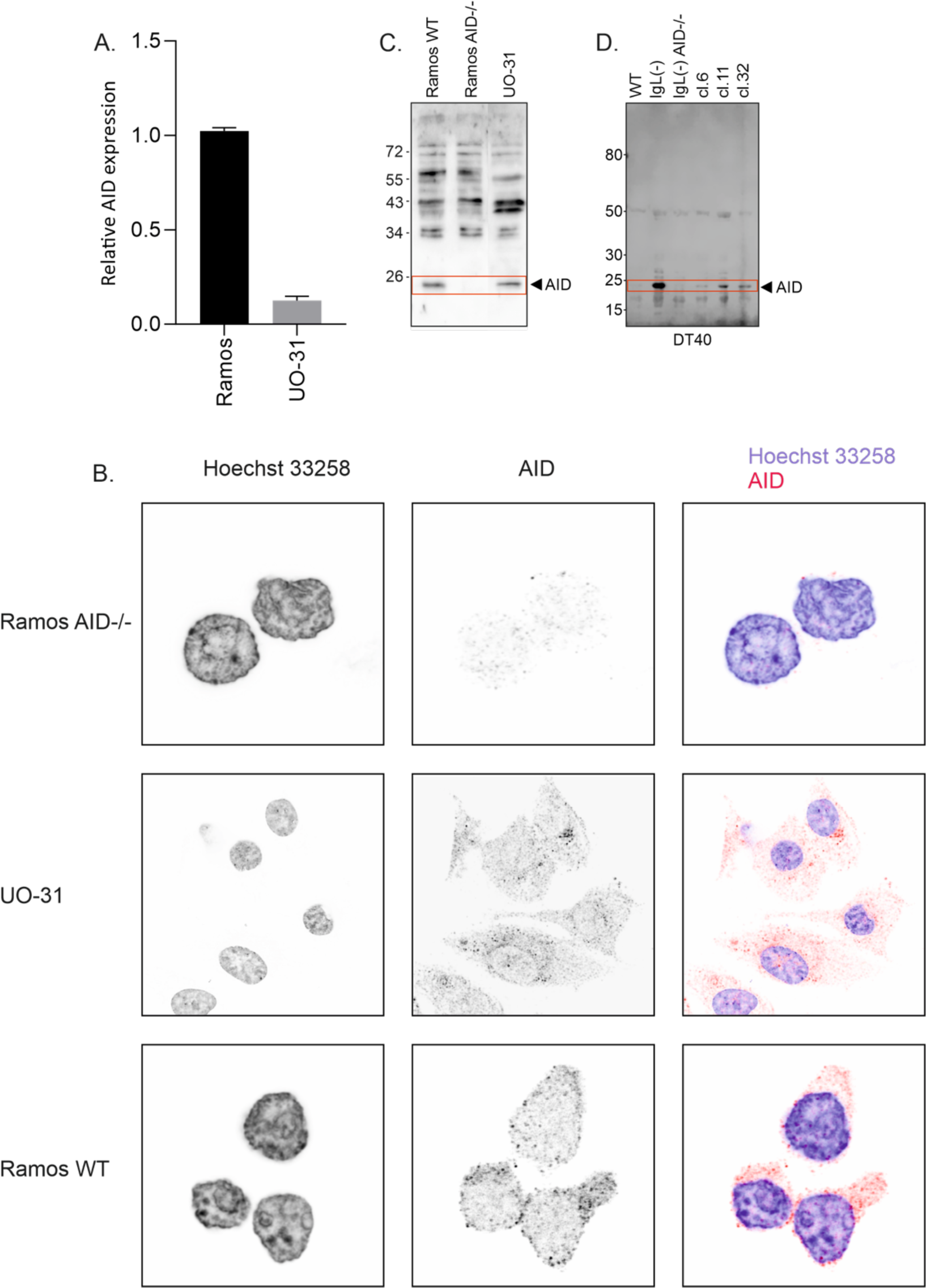
Endogenous AID expression in assayed cell lines. A. Relative AID expression in Ramos WT and UO-31 cell lines measured with RT-qPCR. B. Immunocytochemistry detection of AID in Ramos WT, Ramos AID-/. andUO-31 cell lines. C. Western blot detection of AID in Ramos WT, Ramos AID -/- and UO-31 cell lines. D. Western blot detection of AID in DT40 WT, IgL(-), IgL(-) AID -/- cells and IgL(-) derived clones 6, 11 and 32 where SV40 *LT* expression cassette is integrated to Ig locus.

**Figure S4.**
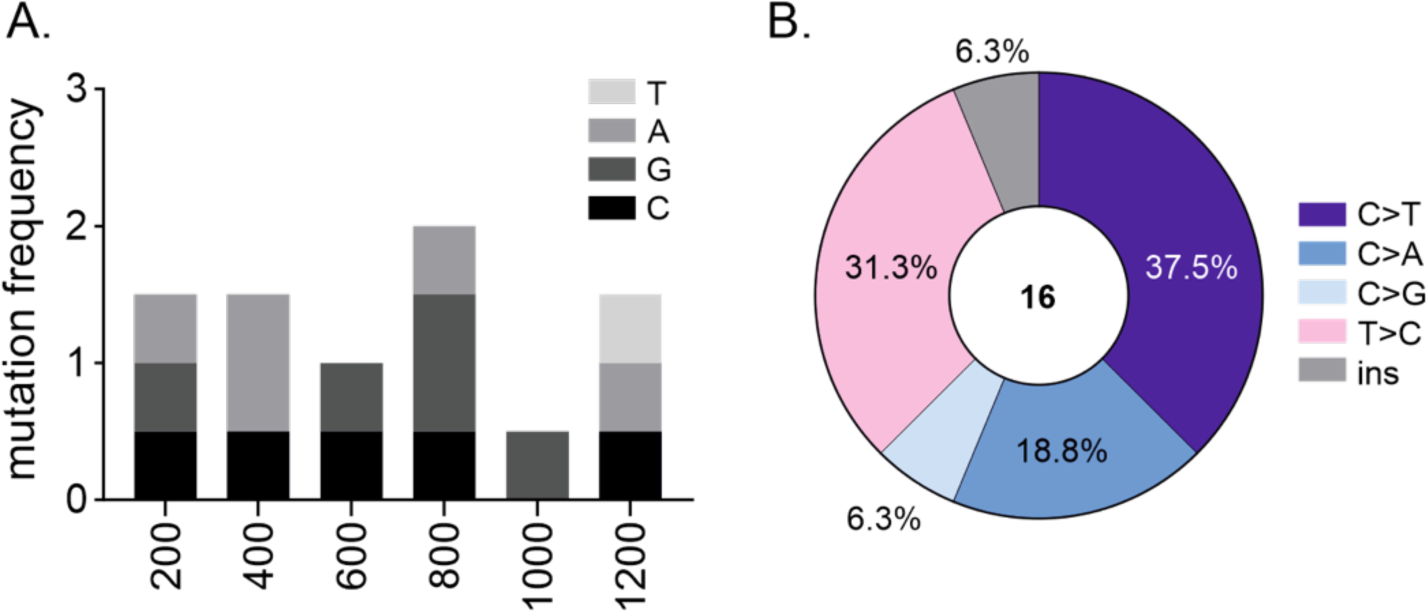
Mutation type and distribution in *GFP2a7a* gene in UO-31 cells. A. Mutation distribution along *GFP2a7a* gene. Mutations targeting different bases are indicated in different colors. Mutation frequency (number of mutations/number of sequences per bin) is shown. B. Mutation types in *GFP2a7a* gene. Substitutions are presented in six substitution classes: C>A, C>T, C>G, T>A, T>C, T>G. In addition, deletions and insertions are shown.

**Figure S5.**
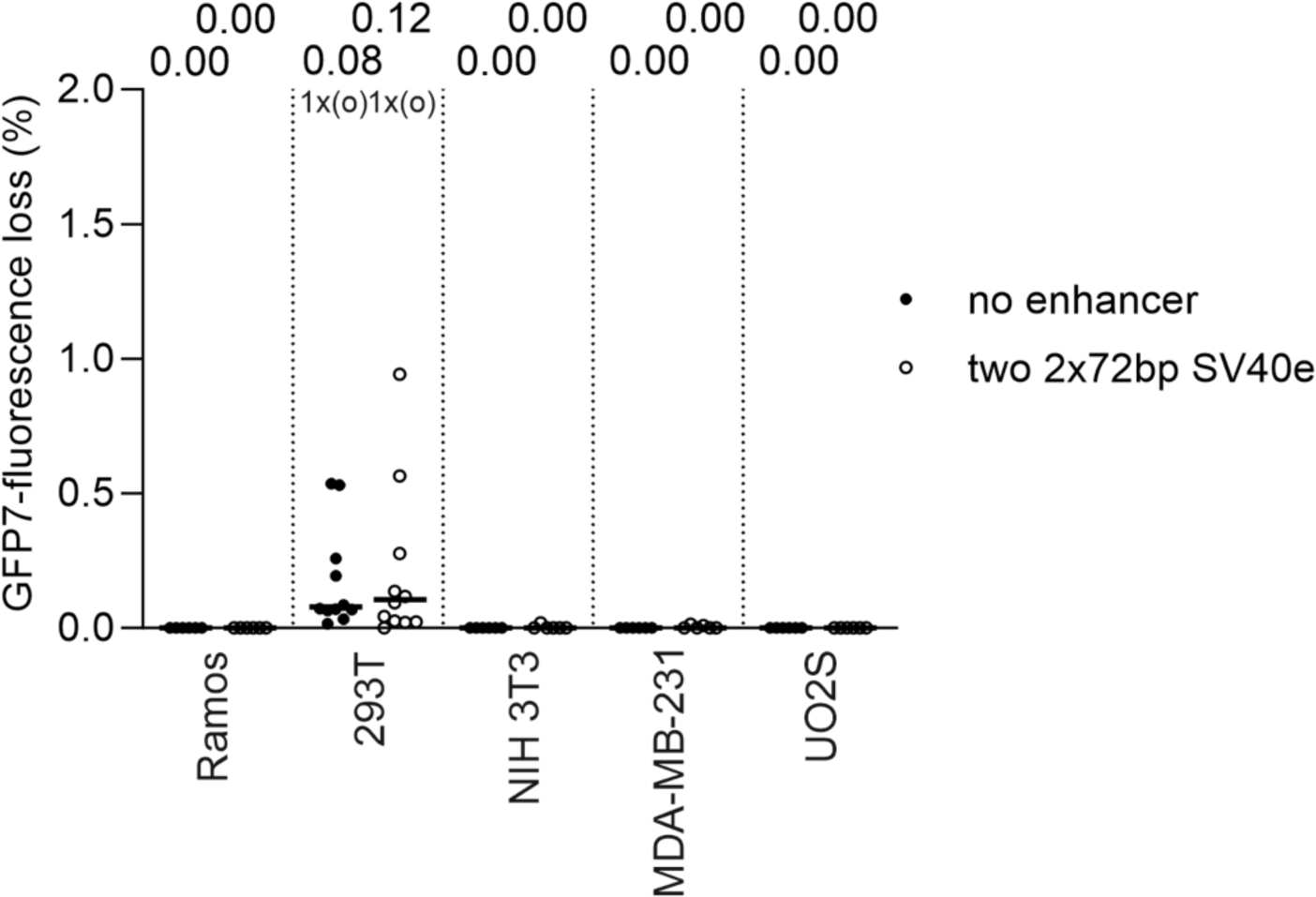
GFP fluorescence loss in non-B cells without AID expression. GFP fluorescence loss of two 2×72bp SV40 enhancer measured in GFP fluorescence loss assay in GFP7 reporter in Ramos AID-/-, 293T, NIH 3T3, MDA-MB-231 and UO2S cell lines which were not transduced with AID-mCherry expression vector. Median values are shown.

**Figure S6.**
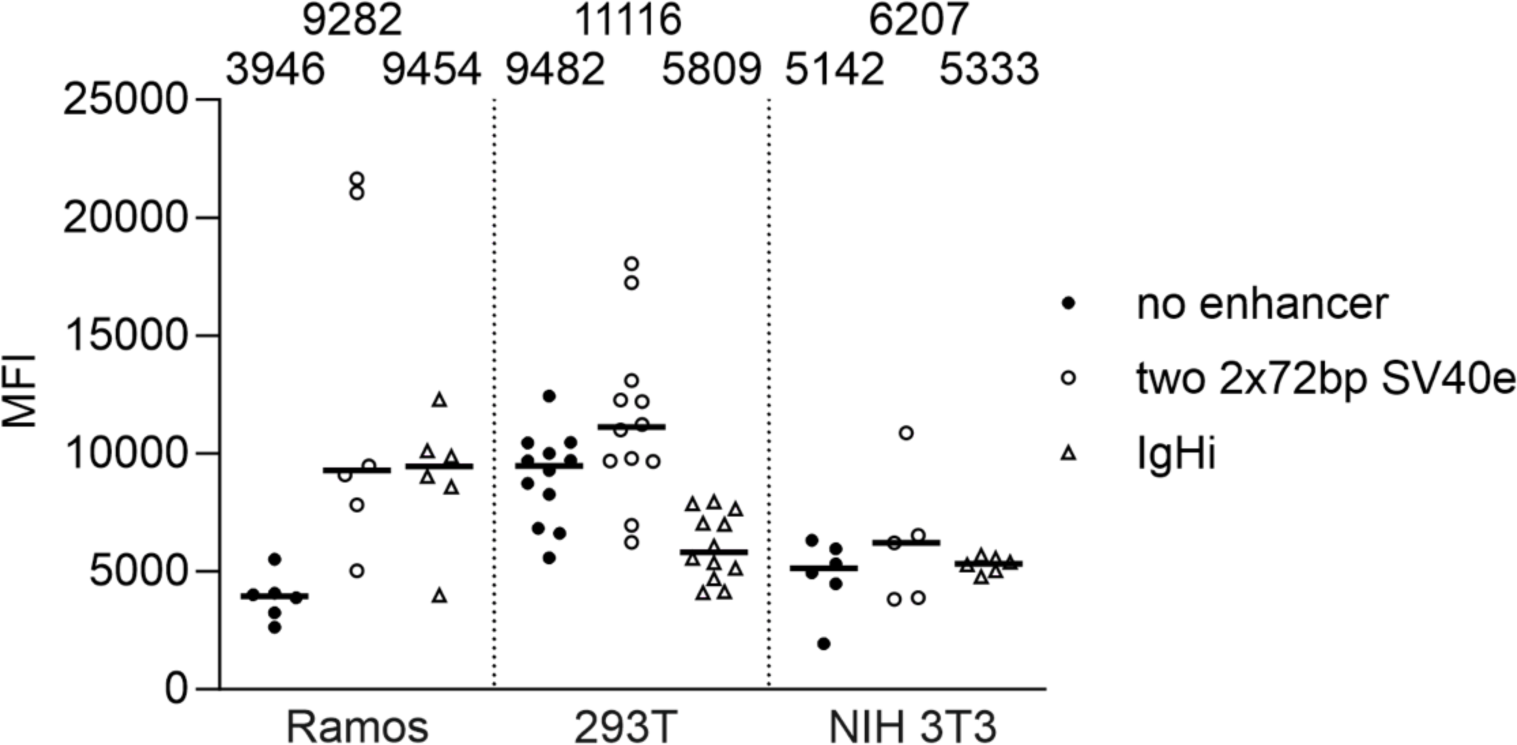
Mean fluorescence intensity of two 2×72bp SV40 enhancer and IgHi. Mean fluorescence intensity of two 2×72bp SV40 enhancer and IgHi measured in GFP fluorescence loss assay in GFP7 reporter in Ramos AID-/-, 293T, NIH 3T3 cell lines.

## Materials and methods

### Cell culture

DT40 cells were cultured at +40 °C, 5% CO2, 90% humidity. Growth media included RPMI 1640 HEPES modification (Sigma) with 10% FBS (HyClone), 1% NCS (Biowest), 1x penicillin-streptomycin antibiotic (Gibco), 1x L-glutamine (Gibco), and 50 µM β-mercaptoethanol. Ramos cells were cultured at +37 °C, 5% CO2, 90% humidity. Growth media included RPMI 1640 HEPES modification (Sigma) with 10% FBS (Gibco), 1x penicillin-streptomycin antibiotic (Gibco) and 1x L-glutamine (Gibco). UO31, H28, 293T, U20S and MDA-MB-231 cells were cultured at +37 °C, 5% CO2, 90% humidity. Growth media included Dulbecco’s Modified Eagle’s Medium – high glucose (Sigma) supplemented with 10% FBS (Gibco) and 1% penicillin-Streptomycin (Sigma).

### Cloning to GFP vectors

SV40 enhancer was amplified with PCR using Q5 High-Fidelity DNA Polymerase (NEB) from plasmid template. Enhancers were cloned into GFP2, GFP4 -and GFP7 expressing vector using In-Fusion cloning kit (Takara) according to manufacturer’s protocol. *Nhe*I/*Spe*I site was used for cloning to GFP2 and GFP4 vectors and HpaI site was used for cloning to GFP7 vector. The mutated SV40 enhancer sequences were created with QuickChange site-directed mutagenesis kit (Stratagene) or using InFusion site-directed mutagenesis. The cloned plasmids were isolated with GeneJET Plasmid Miniprep Kit (Thermo Scientific) or QIAprep Spin Miniprep Kit (QIAGEN). For GFP2 and GFP4 vectors successfully cloned enhancer constructs were further isolated with ZymoPure II Maxiprep kit (Zymo research).

### GFP loss assay (DT40)

Plasmids containing SV40 enhancer and modifications were transfected to chicken B-cell line DT40 (RRID:CVCL_0249) cells with modifications (*UNG*^-/-^*AICDA*^R/puro^). Assay performance is described in detail in (26,33). At the end of the assay cells were analyzed using Accuri C6 cytometer (BD Bioscience) and Novocyte cytometer. Results were further analyzed with FlowJo software (RRID:SCR_008520) and GraphPad Prism 9 software (RRID:SCR_002798).

### GFP loss assay (Ramos and other cell lines)

The GFP loss assay with the GFP7 reporter was done as previously described (32). Plasmids containing SV40 enhancer and modifications were transduced to Ramos B cells, UO31 cells, U20S cells, MDA-MB-231 cells. GFP expression was assessed by analysis with an LSRII cytometer (Becton Dickinson). Results were further analyzed with FlowJo software.

### Luciferase assay

Tested SV40 enhancer and modifications were cloned to SalI/BamHI site at pGL4.23 vector. 20 µg of the plasmid was co-transfected with 2.5µg of pGL4.75 Renilla luciferase control vector (Promega) into DT40 AIDR/puro UNG-/- cells. Transfection was performed using the Amaxa Nucleofector kit V program B-023 (Lonza) or Xcell PBS protocol. The Dual-Glo Luciferase Assay System (Promega) was used to determine the relative activity of firefly luciferase to Renilla luciferase. Assay was performed according to manufacturer’s protocol.

### Statistical analysis

Mann–Whitney U test was performed to evaluate statistical significance of differences in the medians of GFP fluorescence loss and MFI. Unpaired t-test was used to evaluate statistical significance of relative luciferase activity. Fisher’s exact test was used to evaluate statistical significance of WR**<u>C</u>** mutations compared to overall C targeting mutations. Statistical analyzes were performed using GraphPad Prism 9 software.

### *LT* area insertion to DT40 genome

SV40 *LT* area was amplified with PCR using Q5 High-Fidelity DNA Polymerase (NEB). Template was SV40 genome WT4 plasmid which was received as a gift from James DeCaprio. Primers were designed according to In-Fusion primer design protocol. Cloning to vectors was performed with In-Fusion cloning (Takara) kit according to the manufacturer’s protocol. Cloning sites were NheI/SpeI (GFP2).

GFP2 vector containing SV40 *LT* area was transfected to DT40 IgL(-) cells (REF). Transfection was conducted as in GFP loss assay protocol (26). Day after transfection blasticidin selection was added to kill the cells where expression cassette has taken the plasmid in. Sufficient blasticidin concentration (final) is 20 µg/ml. First 15 µg/ml was used and the additional selections were made with 20 µg/ml. After 7-9 days single clones were picked and expression cassette targeting to right locus was confirmed with puromycin selection (final concentration 1 µg/ml). After this, targeted clones were cultured for 12 weeks in order to mutations to accumulate. Culture procedure is described above. Genomic DNA and proteins were extracted at certain time points. Zymo gDNA miniprep kit was used for DNA extraction. Protein extraction is described separately.

*LT* area was amplified from 12-week gDNA with PCR using Q5 High-Fidelity DNA Polymerase (NEB) and cloned to pUC19 vector to BamHI/HindIII site. Primers were designed according to In-Fusion primer design protocol. Cloning was performed with In-Fusion cloning (Takara) kit according to the manufacturer’s protocol. The cloned plasmids were isolated with GeneJET Plasmid Miniprep Kit (Thermo Scientific). Sequencing was performed at Eurofins Genomics, Germany. Sequences were analyzed using SnapGene software (RRID: SCR_015052).

### *LT* area insertion to Ramos and UO-31

SV40 *LT* area was amplified with PCR using Phusion Hot Start II High-Fidelity DNA Polymerase (Thermo Scientific). Template was SV40 genome WT4 plasmid which was received as a gift from James DeCaprio. The PCR product was cloned via In-Fusion cloning system (Takara) into HpaI restriction site in GFP7 vector. Virus production was performed as described previously (Senigl et al., 2019). Ramos WT and UO-31 cell lines were transduced with the vector in multiplicity resulting in 1-2% of GFP-positive cells in culture to achieve one copy of the vector per cell. Three days after the infection, the GFP-positive cells were sorted, expanded and cultured 12 weeks. Genomic DNA and proteins were extracted at the end of culture.

### Protein extraction and western blot analysis in DT40 cells

10M cells in a suspension were centrifuged at 500xg, 5 min at +4°C. Supernatant was discarded and cells were resuspended to 500 µl of PBS and centrifuged at 2000xg, 3 min at +4°C. Supernatant was discarded and cells were resuspended to 100 µl ice-cold RIPA buffer with 1x protease inhibitor cocktail and incubated 1 hour on ice. After this, lysate was centrifuged at 10 000xg, 10 min at +4°C. Supernatant was transferred to clean Eppendorf tube and 33.3 µl of 4x NuPage LDS Sample buffer with 0.2M DTT was added. Lysate was incubated at +70°C for 10 minutes and stored at -80°C.

10 ul of protein lysate and 5 µl of Spectra BR marker (Thermo scientific) was loaded to NuPAge Bis-Tris 4-12% gel with MOPS buffer in Xcell system. Samples were run at 180V for 1 hour. After this, proteins were transferred to nitrocellulose membrane using Xcell blotting system using manufacturers protocol in RT, 30V for 1 hour. Membrane was blocked with 5%-BSA-TBS for 2-3 hours at RT. Primary antibody incubations were performed overnight at +4°C in rocker (LT and AID) or at RT, 15 min in rocker (GAPDH). Secondary antibody incubations were performed at dark, RT, 2-3 hours (LT and AID) or 15 minutes (GAPDH), in a rocker. Membrane was imaged using LICOR Odyssey Fc machine. Membrane was washed between blocking, changing the antibodies and before imaging for 3×5min with TBS.

Primary antibodies used in western blot were αAID 30F12 (Rabbit) Cell Signaling 1:1000 dilution in 5%-BSA-TBS, αSV40-LT pAb419 Genetex (Mouse) 1:100 dilution in 5%-BSA-TBS and αGAPDH Hytest 5G4 (Mouse) 1:10 000 dilution in 5%-BSA-TBS. Secondary antibodies used were IRdye αRabbit 680RD 1:2500 in 5%-BSA-TBS for detecting AID expression, IRdye αMouse 800CW 1:5000 in 5%-BSA-TBS for detecting LT expression and IRdye αMouse 680RD 1:10000 in 5%-BSA-TBS for detecting GAPDH expression.

### Protein extraction and western blot in Ramos B cells and UO31 cells

Cells were washed with ice-cold PBS, the supernatant was removed by aspiration then 1x Laemmli sample buffer (0.5 mL per 5×10^6^ cells/60 mm dish) was added. The cell lysate was boiled for 5min and sheared through a needle 10x.

5ul of protein lysate (Ramos WT and AID -/- cells), 30 ul of (UO31, H28 cells), and 5µl of color protein standard marker (Bio Labs) was loaded to 13% (for AID) or 12% (for LT) SDS-PAGE Gel with Elfo buffer in Xcell system. Samples were run at 100V 10min and them 150V 1h. After this, proteins were transferred to xxx membrane (pre-washed in 100% methanol and incubate in buffer blotting 10min) using trans-blot SD semi dry transfer system (Biorad) in RT, 15V for 30min. The membrane was blocked with 5%-BSA-TBS (for AID detection) or 5% milk (for LT detection) for 1h at RT. Primary antibody incubations were performed overnight at +4°C in rocker. Secondary antibody incubations were performed at dark, RT, 1h. The chemiluminescent Detection Substrate-West Pico Plus (Thermofisher) was used for the detection. The images were obtained using the UVITEC Cambridge imaging system. The membrane was washed between blocking, changing the antibodies, and before imaging for 3×5min with TBS. Used primary antibodies were AID Monoclonal Antibody (ZA001) (Mouse) Thermo Fisher Scientific 1:1000 dilution in 5%-BSA-TBS and αSV40-TAg pAb419 Genetex (Mouse) 1:100 dilution in 5%-BSA-TBS. Anti-Mouse IgG HRP linked antibody (Cell signaling) 1:3000 dilution in 5%-BSA-TBS was used as secondary antibody.

### Immunohistochemistry

Ramos WT, Ramos AID -/- and UO31 were seeded on microscope cover glasses (3×10^5^/well in the 6-well plates with the cover glass with poly-L-Lysine). After overnight incubation, the cells were fixed with 4% paraformaldehyde for 15min; washed 3x with PBS, permeabilized with 0,1% Triton-1% BSA in PBS for 10 min; and washed 3x with PBS. The blocking was performed using 2% BSA-0.15% Glycine-10% FCS in PBS at RT for 45 min; washed 3x with PBS. The slides were incubated for 1h at RT with primary antibody (1:20) Mouse Monoclonal Ab anti-AID (ZA001, Invitrogen), in PBS-0.1% BSA-0.4% Tween20, washed, and incubated with secondary antibodies anti-mouse –Cy3 (1:500) for 1 hour at room temperature. Hoechst 33258 for 15min at RT was added to visualize the cell nucleus, and washed 3x with PBS. Images were taken using a Leica SP8 confocal microscope.

### Leica SP8 confocal microscopy acquisition

Samples were acquired using a Leica SP8 FLIM confocal microscope, equipped with 405 nm, 445nm, 552nm, and white light laser. The 63x oil immersion objective was used (HC PL APO CS2 63x Oil, NA 1.4, WD 0.14 mm, DIC), zoom 4 for Ramos cells and zoom 2 for UO31 cells. Hoechst 550 was detected using the laser 405 and detector PMT1 (420-480), and Cy3 was detected using laser 552 and detector PTM4 (554-620). The acquisition of Hoechst and Cy3 were separate in two different channels to avoid background. The laser intensities were adjusted to avoid bleed-through between channels. Data were collected with a resolution of 1024 × 1024 pixels and step every 0.45um. Images were deconvoluted by Huygens software and the contrast was enhanced using ImageJ software. Deconvoluted images were subsequently exported for analysis using Fiji J.

### Reverse transcription and Quantitative PCR

Different cell lines were analyzed: Ramos WT, UO31, H28, 786-O, U20S, MDA-MB-231, and MDMA231 for AID and GAPDH (gene keeping). The reagent RNAzol® RT was used to isolate total RNA. H2Odd was added to the sample and incubated at RT for 15min, samples were centrifuged at RT at 16 000G for 15min. The supernatant was transferred and 75% ethanol was added to precipitate RNA 10min at RT. The pellets were kept after centrifugation for 10 min at 12 000G at 15°C and washed 2x with ethanol. H2Odd was added to the dry pellet and heated at 55°C for 10min.

DNAase treatment using the kit RQ1 RNase-free DNase (Promega, ref. M6101) was performed. The prepared RNA was reversely transcribed with Protoscript II Reverse Transcriptase (NEB, M0368) using random primers from the kit.

An amount of 2 μL of cDNA was then used for quantitative PCR with the MESA GREEN qPCR MasterMix Plus (Eurogentec) and primers for AID (Forward 5′ AATTCAAAAATGTCCGCTGGGC*T3′, Reverse 5′AGCGGAGGAAGAGCAATTC*C3′) or GAPDH (Forward 5′AGGGCTGCTTTTAACTCTGG*T3′, Reverse 5‘CCCCACTTGATTTTGGAGGG*A3‘).

To generate the standard curve for absolute quantification of gene expression, we used serial dilution of Ramos WT. The samples (in triplicates) were run in a Bio-Rad CFX96^TM^ Real-Time instrument with a 3-step protocol: one cycle of 3 min at 95°C, then 40 cycles of 15 s at 95°C, 20 s at 60°C, and 20 s at 72°C and final polymerization at 72°C for 10 min. Cycles of quantification (Cq) values were generated by the CFX Manager software.

## Acknowledgements

This work was supported by the Czech Science Foundation (project 22-30384S to F.Š.) and Czech Academy of Sciences (Premium Academiae Award 2018 to J.H.), the Finnish Cultural Foundation, the Sigrid Juselius Foundation, the Turku University Foundation (to J.A.), the Finnish Cultural Foundation Kymenlaakso Regional Fund, the Alfred Kordelin Foundation (to A.S.) and Cancer Society of Southwest Finland (A.S. and J.A.), NIH R01 AI 127642 (D.G.S.). FŠ and J.H. were supported by the project National Institute of Virology and Bacteriology (program EXCELES, no.LX22NPO5103) funded by the European Union–Next Generation EU; we also acknowledge institutional support from the project RVO (68378050).

